# Characterization and visualization of global metabolomic responses of *Brachypodium distachyon* to environmental changes

**DOI:** 10.1101/2022.05.11.491395

**Authors:** Elizabeth H. Mahood, Alexandra A. Bennett, Karyn Komatsu, Lars H. Kruse, Vincent Lau, Maryam Rahmati Ishka, Yulin Jiang, Armando Bravo, Benjamin P. Bowen, Katherine Louie, Maria J. Harrison, Nicholas J. Provart, Olena K. Vatamaniuk, Gaurav D. Moghe

**Affiliations:** Plant Biology Section, School of Integrative Plant Science, Cornell University, Ithaca, NY, USA; Institute of Analytical Chemistry, Department of Chemistry, Universität Für Bodenkultur Wien, Vienna, Austria; Department of Cell and Systems Biology, University of Toronto, Canada; Michael Smith Laboratories, University of British Columbia, Vancouver, Canada; Boyce Thompson Institute, Ithaca, NY, USA; Soil and Crop Sciences Section, School of Integrative Plant Science, Cornell University, Ithaca, NY, USA; Donald Danforth Plant Science Center, Olivette, MO, USA; Environmental Genomics and Systems Biology Division, Lawrence Berkeley National Laboratory, Berkeley, CA, USA; Department of Energy Joint Genome Institute, Lawrence Berkeley National Laboratory, Berkeley, CA, USA

**Keywords:** Computational biology, Metabolomics, Mass spectrometry, Abiotic stress, Mycorrhizal symbiosis, Brachypodium

## Abstract

Plant responses to environmental change are mediated via changes in cellular metabolomes. However, <5% of signals obtained from tandem liquid chromatography mass spectrometry (LC-MS/MS) can be identified, limiting our understanding of how different metabolite classes change under biotic/abiotic stress. To address this challenge, we performed untargeted LC-MS/MS of leaves, roots and other organs of *Brachypodium distachyon*, a model Poaceae species, under 17 different organ-condition combinations, including copper deficiency, heat stress, low phosphate and arbuscular mycorrhizal symbiosis (AMS). We used a combination of information theory-based metrics and machine learning-based identification of metabolite structural classes to assess metabolomic changes. Both leaf and root metabolomes were significantly affected by the growth medium. Leaf metabolomes were more diverse than root metabolomes, but the latter were more specialized and more responsive to environmental change. We also found that one week of copper deficiency shielded the root metabolome, but not the leaf metabolome, from perturbation due to heat stress. Using a recently published deep learning based method for metabolite class predictions, we analyzed the responsiveness of each metabolite class to environmental change, which revealed significant perturbations of various lipid classes and phenylpropanoids such as cinnamic acids and flavonoids. Co-accumulation analysis further identified condition-specific metabolic biomarkers. Finally, to make these results publicly accessible, we developed a novel visualization platform on the Bioanalytical Resource website, where significantly perturbed metabolic classes can be readily visualized. Overall, our study illustrates how emerging chemoinformatic methods can be applied to reveal novel insights into the dynamic plant metabolome and plant stress adaptation.

## Introduction

The central dogma of molecular biology extends from genes to transcripts to proteins. These proteins, however, exert an effect on the phenotype eventually through altering metabolites. Agronomically important traits such as yield, nutritional quality, flavor characteristics and stress response are all controlled by underlying metabolic pathways. A revolution in sequencing over the past decade has provided unparalleled insights into the transcriptomic and epigenomic perturbations due to genotypic and environmental changes, yet the global metabolome largely remains a black box, primarily due to our inability to identify compounds from metabolomics data (Chaleckis et al., 2019; Salem et al., 2020). It is estimated that over a million compounds are produced across the plant kingdom (Afendi et al., 2012), with individual plants producing thousands of metabolites (Fernie, 2007). However, <5% of these signals can be annotated using spectral matching (da Silva and Dorrestein, 2015). Thus, patterns of global metabolomic changes still remain unknown despite the importance metabolites have to plant fitness and human society.

To assess metabolomic changes due to genetic variation, developmental progression and environmental changes, gas chromatography mass spectrometry (GC-MS) and liquid chromatography mass spectrometry (LC-MS) remain the workhorse approaches, with LC-MS typically detecting a much broader set of the metabolome. Although diverse algorithmic innovations have aided in metabolome assessments (Brouard et al., 2016; Tsugawa et al., 2016; Schymanski et al., 2017; Dührkop et al., 2019), LC-MS peaks are primarily annotated using MS/MS spectral matching with entries from public databases (Horai et al., 2010; Wang et al., 2016; Guijas et al., 2018). While correct predictions are indeed obtained in this manner, plant-derived compounds are underrepresented in public databases (Fukushima and Kusano, 2013; Shahaf et al., 2016), which potentially produces false positives in the limited numbers of compounds identified. Partly due to this limitation, many LC-MS based studies are targeted or semi-targeted, and end up analyzing a small but identifiable portion of the metabolome (Itkin et al., 2013; Okazaki et al., 2013; Bromke et al., 2015; Tohge et al., 2016; Šimura et al., 2018). This strategy produces robust insights, but global shifts in the metabolome and their genetic drivers cannot be assessed via targeted studies. Identifying such patterns can provide novel insights into metabolic plasticity and plant responses to stress conditions, which are important for addressing challenges of agricultural productivity due to climate change, overpopulation and degrading soil quality.

In recent years, two important resources have emerged for the analysis of global untargeted tandem LC-MS (LC-MS/MS) data. Firstly, the machine learning (ML) based tool CANOPUS (Dührkop et al., 2021) enables prediction of metabolite structural classes based on the MS/MS spectrum, providing novel insights into the metabolome composition. For example, even if specific compounds are not identified, recognizing that “flavonoids” increase in abundance under UV stress provides significant biological insights into the plant’s stress response. Secondly, independent of compound annotation, approaches adapted from information theory can inform about the gross and/or specific shifts in plant metabolomes (Li et al., 2020; Zu et al., 2020). In this study, we combine these two approaches to illuminate global changes in plant metabolomes under different conditions.

Specifically, we assessed the metabolome of *Brachypodium distachyon* (Brachypodium) under different conditions **(Fig. 1)**. Brachypodium is a model C3 grass species in the Poaceae family that shared a common ancestor with rice (*Oryza sativa*) ~50 million years ago and Triticeae (wheat, barley) ~35 million years ago (Charles et al., 2009). The short stature of Brachypodium and its fast growth cycle make the species a convenient model for understanding not only Poaceae biology but also for biofuel research (Brkljacic et al., 2011; Douché et al., 2013; Marriott et al., 2014; Le Bris et al., 2019). The main goals of this study were to: (i) assess Brachypodium metabolome reconfigurations across different organs and a breadth of environmental conditions, (ii) identify the metabolite classes most perturbed by different stresses, (iii) discover condition-specific metabolites that may serve as stress biomarkers, and (iv) establish a platform for visualization of the global metabolome changes. Towards these goals, we first performed LC-MS/MS from 17 different organ-condition combinations, including agriculturally relevant conditions such as copper deficiency, heat stress, low phosphate, and arbuscular mycorrhizal symbiosis. We used CANOPUS and information theory derived metrics to compare control vs. test metabolomes across different organs, and characterize additional metabolome changes through co-accumulation modules and biomarker detection. Finally, these changes were visualized using a novel representation on the Bio-Analytic Resource for Plant Biology (BAR) website. Overall, our findings provide new insights on the global and more specific metabolic perturbations in Brachypodium under different conditions.

**Figure 1:**
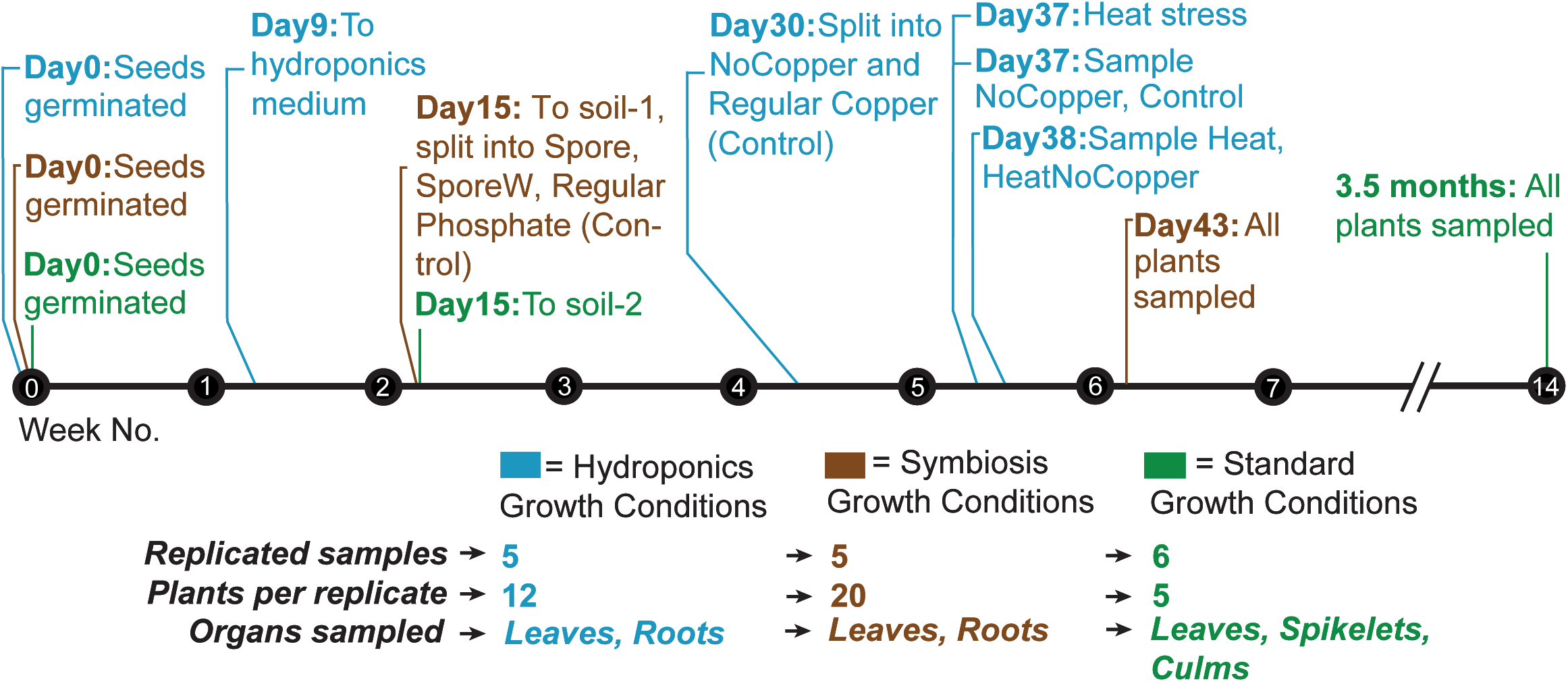
Timeline and Schematic of the Experimental Design. The number of samples, plants per replicate, and organs sampled for each set of growth conditions is shown, along with the timeline of important events such as treatment induction and harvesting. Divergent growth and stress conditions were chosen to induce variability in metabolic profiles. Days are counted post germination. Soil-1 and soil-2 refer to different soil mixes. The germination protocol for Hydroponics seeds was distinct from the germination protocol for the other growth conditions (see Supplemental Methods).

## Results

### Experimental design and pre-processing of metabolome data

Brachypodium plants were grown to different ages and under different growth conditions in order to produce significant metabolome perturbations. Roots, leaves (young and mature combined), and in some cases, culms and spikelets were sampled. Overall, 17 organ-condition combinations were sampled, with plants grown across three major regimens: Hydroponics (Hydro), Symbiosis (Sym), and Tissue (Tis) **(Fig. 1, Supp. Fig. 1)**. Hydro treatments consisted of regular Cu (Control), Cu deficiency (NoCopper), heat stress (Heat), and heat stress under Cu deficiency (HeatNoCopper). The Symbiosis treatments consisted of plants grown with regular amounts of phosphate fertilization (Control), low phosphate treated plants inoculated with a solution of *Rhizophagus irregularis* spore growth medium (SporeW, not containing any spores i.e. mock treatment) and low phosphate treated plants inoculated with *R. irregularis* spores (Spore). The Tissue regimen involved growing plants in regular soil until maturity. The effectiveness of the copper deficiency treatment and presence of colonization were verified through semi-quantitative RT-PCR of copper deficiency and fungal symbiosis marker genes, respectively (**Supp. Fig 2**). All samples were analyzed via LC-MS/MS in both positive and negative mode to obtain a comprehensive, quantitative snapshot of their metabolome.

After peak deconvolution and alignment, metabolite values were filtered using a sequence of steps **(Supp. Figs. 3,4)**. To enable comparisons between different LC-MS runs, we first tested five different data normalization approaches **(Supp. File 1)** and selected Variance Stabilized Normalization (VSN) as the most appropriate based on performance as well as availability of the algorithm **(Supp. Table 1; Supp. File 1)**. Data imputation was also performed to fill in values lost due to Orbitrap LC-MS detection limits. To ensure that either step does not alter the overall underlying data structure, we first determined the effect of performing imputation before vs. after normalization using a dummy dataset where actual peak areas were randomly replaced by zeros. The degree of error in normalization-imputation and imputation-normalization was quantified. Overall, both normalization orders had almost identical errors **(Supp. Fig. 5)**. Thus, given precedence (Mock et al., 2018; Chong et al., 2019), we first imputed peak areas using k-Nearest Neighbor and normalized the imputed areas using VSN for further downstream analyses.

VSN maximized correlations among replicates while maintaining low correlations between different treatment groups **(Supp. File 2)**. The above ground tissues were found to have more peaks as well as a higher total peak abundance than the roots **(Supp. Fig 6; Supp. File 3)**. The largest number of metabolite signals in both organs were observed in Sym samples, indicating that growth media also influenced the Brachypodium metabolome. The high numbers of peaks seen in the Sym Spore root samples may include metabolites of fungal origin. Correlations between leaf vs. root, and between control vs. treatment, were respectively much or slightly lower than among replicates **(Supp. File 2)**, putatively identifying two other axes of metabolomic divergence between samples. To investigate these further, we first performed a global assessment of similarities and differences between the metabolomes under different conditions.

### The root metabolome is less diverse but more specialized and more stress-inducible than the leaf metabolome

Using the normalized, imputed datasets, we quantified the impact of each stress on the root and leaf metabolomes. As expected, Principal Components Analysis (PCA) identified the organs and the growth media as stronger drivers of metabolic variation in our samples than the stresses. While PC1, explaining 46.88% and 45.6% of the metabolic variation between samples (in positive and negative mode, respectively) was indicative of organ-wise differences, PC2 (12.97%, 13.77%) revealed a substantial impact of the growth medium (soil type, hydroponics) on the root and leaf metabolomes (**Supp. Fig. 7**). PCA as well hierarchical clustering (**Supp. Figs. 8,9**) validated close clustering of replicate samples as well as highlighted set-wise impact of stresses. For the Hydro set, NoCopper (copper deficiency) was clustered with Control in both leaves and roots, while for the Sym set, SporeW was the more impactful condition for leaves and Spore for the roots. HeatNoCopper clustered closer to Heat than NoCopper in both roots and leaves, indicating that the majority of metabolomic differences in this combined stress was due to Heat. When PCAs were differentiated by organs (**Supp. Fig. 7 B,C,E,F**), the effect of different stresses could be observed. Overall, the leaf metabolomes were less impacted by the stresses than root metabolomes.

To further quantify the impact of each stress on the overall sampled metabolome, we used three information-theory based measures – Diversity (H), Specialization (δ, measuring uniqueness/differentiation) and Relative Distance Plasticity Index (RDPI, measuring overall perturbation including up and down-accumulation). We first assessed the metabolome differences in non-stress conditions. More peaks as well as more uniformity in the peak areas can increase Diversity; thus, given leaves consistently have more peaks than roots, culms, and spikelets (**Supp. Fig. 6**), their Diversity is the highest (**Fig. 2A,B; Supp. Fig. 10A,C**). However, roots and spikelets are more metabolically specialized. The degree of specialization and to some extent, Diversity, were clearly dependent on the growth medium and stress (**Fig. 2A,B; Supp. Fig. 10**). Roots were more specialized in the hydroponic medium (except Sym Spore root) but leaf metabolome was more specialized in soil (**Supp. Fig. 10B,D**). Intriguingly, the observation of spikelets being metabolically specialized is congruent with a similar observation in the *Nicotiana attenuata* anthers (Li et al., 2016b), indicating that the metabolic uniqueness of the reproductive tissues may be a conserved trait across monocots and dicots.

**Figure 2:**
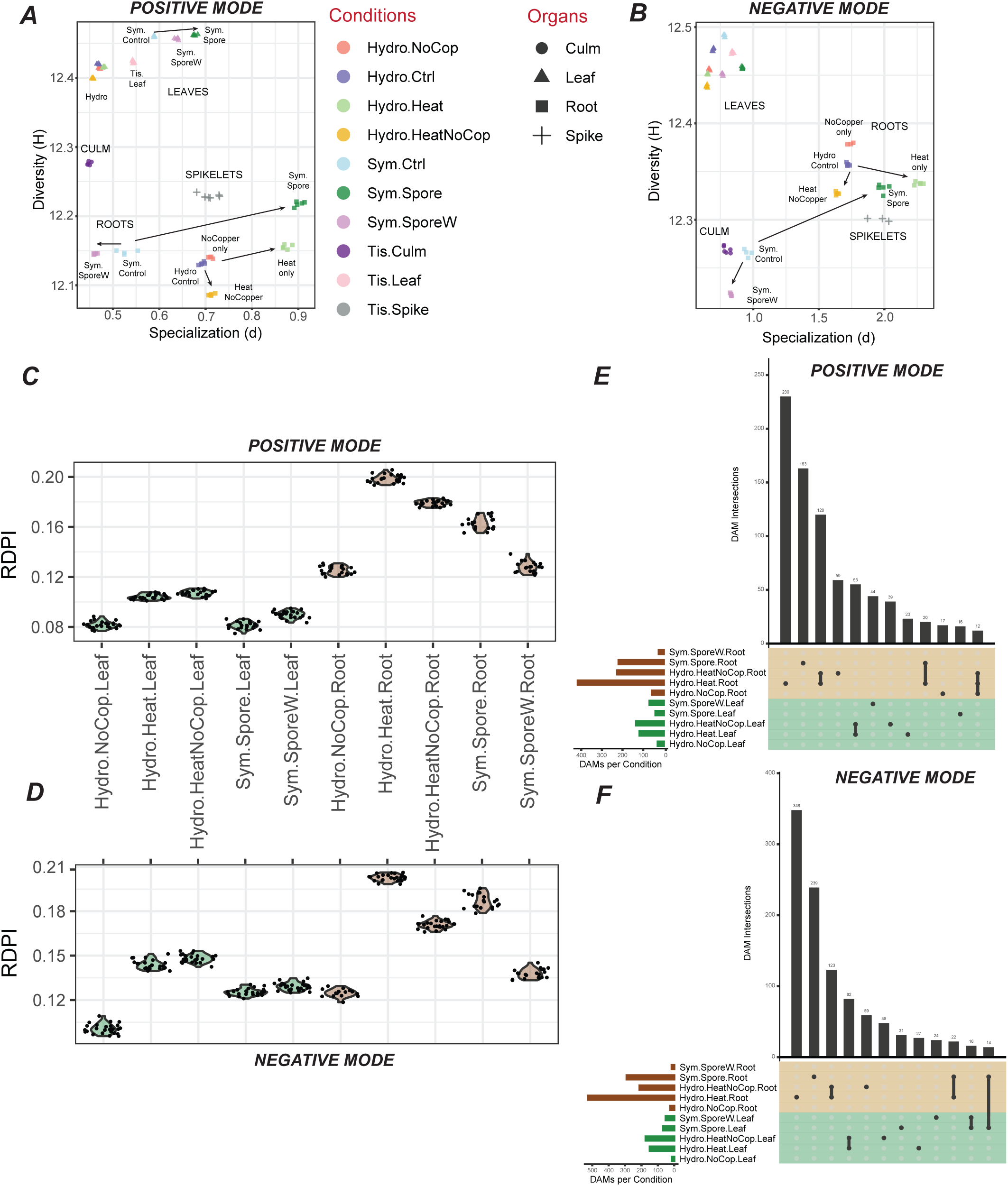
Comparison of metabolomic perturbations among conditions. (A), (B) Diversity vs. Specialization per condition, with organs depicted as different shapes and conditions as different colors. Annotations are added onto these plots for ease of interpretation. (C), (D) RDPI per stress condition. (E), (F) Upset plots of Differentially Abundant Peaks (DAP) per stress condition, inclusive of up- and down-accumulated peaks. Intersections (vertical bars) depict the number of DAPs in common to sets of conditions. Only sets with at least 10 peaks are shown. (A), (C), (E) positive mode (B), (D), (F) negative mode.

Although differences in specialization and diversity among leaf metabolomes were low, many stresses elicited statistically significant changes (Kolmogorov Smirnov [KS] test, **Supp. Table 2**). Overall, the stresses appeared to disrupt foliar metabolism far less than that of the roots – especially for leaves from hydroponically grown plants – as indicated by tight clustering of leaf stresses with their controls. In positive mode **(Fig. 2A)**, specialization cleanly separated out leaf samples into their growth conditions, but this was not seen in negative mode **(Fig. 2B)**, and in both ionization modes, leaf samples had relatively low specialization. Taken together with the relatively low RDPI values observed for leaf samples (**Fig. 2C,D**), these results indicate that the leaf metabolome is more robust/less responsive to temporary environmental changes than the root metabolome.

In contrast, the specialization and RDPI of roots were significantly influenced by stress. In both ionization modes, we found that roots had higher RDPI (i.e. greater metabolome perturbation) than leaves (except for SporeW, in which leaves had similar RDPIs to roots in negative mode) (**Fig. 2C,D**). Hydro roots had a higher baseline (Control) specialization than Sym (**Supp. Fig. 10B,D**), indicating the presence of hydroponics-specific peaks. However, in both ionization modes, Heat roots and Spore roots had the highest specialization and RDPI. Specialization is a sum of the “degree of specificity” of each metabolite signal across the different conditions, thus, high specialization in Heat and Spore indicates a greater representation of metabolites that are uniquely changing under these conditions alone. Interestingly, specialization of the HeatNoCopper roots was similar to Control roots **(Fig. 2A,B)**, while its RDPI was intermediate between NoCopper and Heat **(Fig. 2C,D)**. These observations suggest that the impact of heat stress on the global root metabolome was less drastic under copper deficiency, which is contradictory to our expectation that HeatNoCopper roots would show a greater perturbation than Heat roots given a combination of two stresses.

To obtain a more granular understanding of the overall induced metabolites, differentially accumulated peaks (DAPs) were estimated in each condition based on FDR-corrected p-values and fold-change criteria (see **Supplementary Methods**; **Supp. File 4**). The pattern of differential accumulation was similar between positive and negative modes (**Fig. 2E,F**). We found that HeatNoCopper and Heat had a high number of DAPs primarily in the roots (**Fig. 2E,F; Supp. Figs. 11,12,13**). Over 200 metabolites were also perturbed under AMS in positive as well as negative mode, however, many of these metabolites could be of fungal origin. Heat and Spore roots had both the highest numbers of DAPs, and unique DAPs, consistent with the finding that they have high RDPI and the highest specialization.

### Assessment of the deep learning-based tool CANOPUS for structural annotation of LC-MS peaks

The above analyses revealed global patterns of change in the Brachypodium metabolome under environmental change. We next sought to understand shifts in specific metabolite classes. While untargeted LC-MS is the method of choice for detecting a diverse range of metabolites, identifying these peaks is a major challenge. We employed two different approaches for annotating the peaks: 1) MS/MS spectral matching using public repositories, and 2) database-free prediction of structure-based metabolite classes using the deep-learning based CANOPUS package in the SIRIUS software (Dührkop et al., 2021). CANOPUS classifies compounds into the multilabel and hierarchical ChemOnt ontology (Djoumbou Feunang et al., 2016), which is similar to the Gene Ontology (GO) for genes (The Gene Ontology Consortium, 2019). As ChemOnt is multilabel, peaks may receive multiple annotations at each level, however, the classifications we report are of each peak’s largest substructure.

Of the 3582 and 2996 singly charged fragmented peaks in positive and negative mode, 2931 (82%) and 2409 (80%) were annotated by CANOPUS at the Superclass level with posterior probability ≥0.5 **(Supp. Fig. 14)**. Of the 26 Superclasses existing for Organic Compounds, 14 and 12 were represented in the positive and negative mode data, respectively **(Supp. Files 5,6)** with Lipids and lipid-like molecules having the most peaks in both ionization modes. To assess the accuracy of these annotations, we identified peaks via public database searches and compared their ChemOnt classes to CANOPUS’ predictions (**Table 1, Supp. File 7**). At each level, we calculated misannotations as the percent of peaks identified using spectral matches that were not given the same annotation by CANOPUS. At the Superclass level, we observed good correspondence between CANOPUS classifications and database identifications in both modes. The median CANOPUS misannotation rates at the Class level, when considering correct Classes as determined by ClassyFire, were 54.4% and 28.2% in positive and negative mode, respectively, indicating that overall CANOPUS predicted Classes well for negative mode only. In positive mode, the most frequently misannotated Classes were Glycerophospholipids (GPs, 65.57% of CANOPUS-predicted GPs were misannotated) and Phenols (73.68% misannotated), although most of the misannotations were within the same Superclass (70% and 50%, respectively). The decrease in agreement between positive mode Superclasses and Classes is largely due to the high misclassification rate of GPs and their high presence (24%) in the identified positive mode compounds.

**Table 1:** Correspondence between peaks identified using spectral matches and their class predictions using CANOPUS.

|  | Positive |  | Negative |  |
| --- | --- | --- | --- | --- |
|  | Identified<sup>1</sup> and Annotated<sup>2</sup> Peaks | % Match<sup>3</sup> | Identified<sup>1</sup> and Annotated<sup>2</sup> Peaks | % Match<sup>3</sup> |
| **Superclass** | 253 | 77.87 | 153 | 77.78 |
| **Class** | 245 | 54.69 | 152 | 70.39 |
| **Subclass** | 215 | 53.49 | 143 | 71.33 |
| **Level 5** | 123 | 56.10 | 120 | 73.33 |

<sup>1</sup> Identified using spectral matches with public repositories
2 Annotated using CANOPUS
3 Percentage of the Identified and Annotated Peaks with matching annotations

We further observed that when discrepancies occurred, it was often due to CANOPUS labeling compounds based upon substructures that are not representative of the whole compound, e.g. labeling Flavonoids as Benzenoids/Hydroxycinnamic Acid and Derivatives, or 1-Palmitoylglycerol as a Fatty Acyl instead of a Glycerolipid. Most misannotated GPs were classified as Fatty Acyls (subclass: linoleic acid and derivates), Sphingolipids (subclass: phosphosphingolipids) or Organonitrogen Compounds (subclass: phosphocholines), suggesting that despite misclassification, CANOPUS was identifying common substructures from MS/MS data. It is important to also note, however, that in instances of disagreement, the specific compound identifications based on spectral matching may be incorrect, and despite that, both methods generally agree on the annotations of substructures of the detected peaks.

In order to further assess the general accuracy of identifications and CANOPUS annotations, and the disagreements between them for GPs, we used MS/MS molecular networking as a complementary approach to cluster compounds with similar fragmentation patterns. We then mapped identifications and CANOPUS Superclasses onto this network (**Fig 3, Supp. Files 8, 9**). We found that some CANOPUS Superclasses tended to form tight sub-networks e.g. 236 out of the 240 CANOPUS-annotated GPs in the negative mode network were clustered together (**Supp. File 9**), along with all of the database-identified GPs. In the positive mode network, we observed two clusters for GPs --one for peaks identified as Glycerophosphocholines/Glycerophosphoserines and another for peaks identified as Glycerophosphoethanolamines (Subnetworks 1 and 2, respectively, in **Fig 3, Supp. File 8**). For other sub-networks (3,4,5 **Fig. 3**), there was good agreement between CANOPUS and identified compound class predictions (**Supp. File 7**). These results suggest that while there is some disagreement between spectral matching and CANOPUS, both methods are reflective of actual molecular substructures. While Class level interpretation is appropriate for peaks in negative mode, Superclass level interpretation is appropriate for positive mode. Thus, while we conduct analyses below using the more specific Class-level annotations, we primarily interpret results from negative mode data.

**Figure 3:**
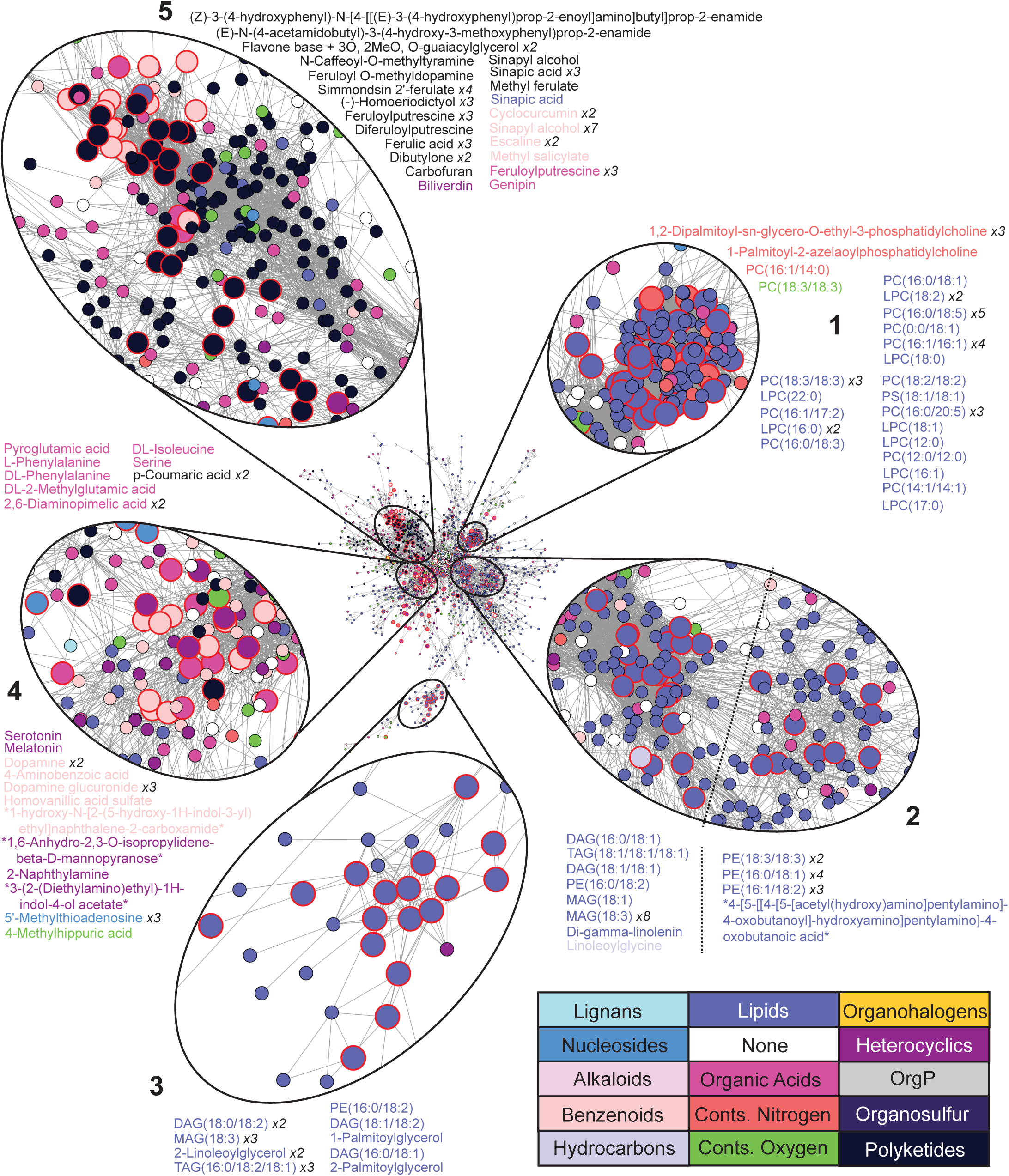
Molecular Networking of peaks in positive mode. Network nodes represent peaks detected in positive mode (in any condition/organ), and edges conntect nodes that have a pairwise cosine score of >70. Large nodes with a red border signify identified peaks. Nodes and identifications (text) are colored with their CANOPUS-annotated Superclass. The number of times each identification occurs in a sub-network is indicated in italics. Asterisks (*) denote an identification spanning multiple lines. The dashed line in subnetwork 2 separates the majority-Glycerolipid section of the subnetwork from the majority-Phosphoethanolamine section. PC = Phosphocholine, L = Lyso, DAG = Diacylglycerol, TAG = Triacylglycerol, PE = Phosphoethanolamine, MAG = Monoacylglycerol.

### Compound class annotation reveals an important role of lipids in the induced stress response

After validating CANOPUS annotations, we sought to determine how different chemical classes were perturbed under the applied stresses, and whether the relevance of a class to a stress or organ could be quantified. The RDPI metric summarizes both up and down regulation of all metabolites in a given class, and thus, is a useful metric to assess a class’ overall perturbation in a given stress **(Supp. Figs. 15, 16; Supp. File 10)**. As expected, RDPI distributions of most Classes (e.g. Organooxygen compounds) were similar to those of the overall metabolome --with roots appearing more inducible than leaves, and Heat, HeatNoCopper and Spore treatments eliciting the largest metabolome changes. However, some Classes – primarily lipids such as Fatty Acyls, Glycerolipids, Glycerophospholipids (GPs), sphingolipids and steroids – deviated from this overall trend. Although the RDPI is a useful metric for quantifying gross metabolomic changes, information on whether peaks are accumulated or depleted under stress conditions is lost. Another issue is that our criteria for calling DAPs is stringent, thus high RDPI does not necessarily translate to more DAPs. Lastly, the RDPI metric for a Class with 1000 metabolites vs. 10 metabolites can appear the same, confounding the true extent of a metabolite Class’ importance in a condition. To address these issues, we identified Classes that were, on average, highly accumulated or depleted in a stress (see Methods), and plotted the abundance changes of individual peaks in those Classes (**Fig. 4**, and **Supp. Fig. 17A**). Many Classes had expected changes in abundance, which corroborates this methodology. For example, in spore-treated samples, lipid Classes such as glycerolipids and GPs decreased (leaves) while prenol lipids and sphingolipids increased (roots) **(Fig. 4)**, consistent with their importance in membrane remodeling and signaling during plant-AMF interactions (Wewer et al., 2014; Macabuhay et al., 2022). Interestingly, more sphingolipids showed a decrease in the leaves, but the pattern was reversed in the roots. Phenolic compounds (Phenols) are known to be induced in the leaves of multiple species under symbiosis (Schweiger and Müller, 2015), which was also observed. In both leaves and roots of Cu-deficient plants, GPs showed both up and down regulation, while sphingolipids were mostly upregulated in the roots. Increase in sphingolipids was also seen in Heat stress. A previous study showed that perturbation of sphingolipid biosynthesis in the roots influences the leaf ionome (Chao et al., 2011), and thus, sphingolipids may play consequential roles in both Cu-deficiency and heat stress. Some lignans were also found to be downregulated in Cu-deficient and heat treated plants in both leaves and roots, consistent with previous observations of lignin biosynthesis affected under copper deficiency (Schulten and Krämer, 2018; Rahmati Ishka and Vatamaniuk, 2020).

**Figure 4:**
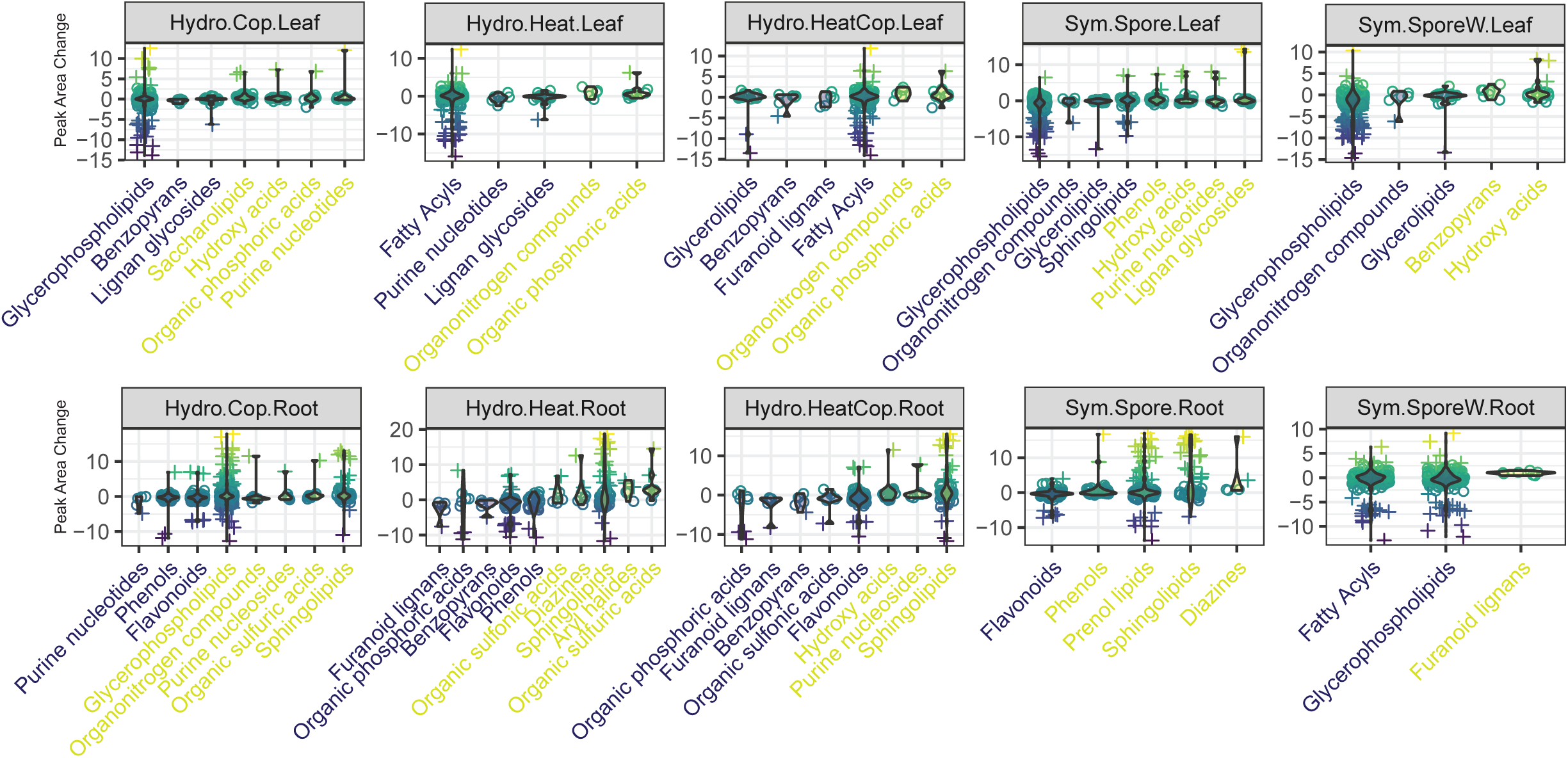
Charting Stress-Induced Shifts of Molecular Classes, Negative Mode. (A) Abundance changes of peaks in response to stress. Each stress depicts Classes that were the most accumulated (Class name in yellow) or diminished (Class name in purple), on average. For a Class to be plotted, its average value must be greater than the 70th percentile or lower than the 30th percentile of all stress-induced peak area changes. Individual metabolites are plotted as circles, outliers are shown as +.

Other classes showed unexpected changes. Although flavonoids are antioxidants, they were, on average, depleted in the roots under multiple stresses (**Fig. 4**). Multiple classes possess outliers present on both sides of the distribution, e.g. Sphingolipids in Spore leaves, suggesting that peaks within the same structural class are not necessarily co-regulated. Finally, we identified classes that were strongly up- or down-accumulated (multiple peaks with area changes > 5 in magnitude) in response to multiple stresses, most of which were lipids e.g. GPs, Fatty Acyls, Sphingolipids and Prenol lipids (**Fig. 4**). These observations suggest that the lipidome is the most stress-responsive portion of the metabolome, possibly resulting from changes in cellular membranes and signaling pathways.

### Co-accumulated peaks have diverse structural classes, and peaks within a class rarely co-accumulate

As many classes showed broad changes in response to a stress, we next assessed the diversity of structural classes among groups of correlated peaks as determined using Weighted Gene Coexpression Network Analysis (WGCNA) (Langfelder and Horvath, 2008) (**Fig. 5A** and **Supp. Figs. 17,18,19**). WGCNA provides a complimentary approach to assign functional hypotheses to metabolite classes under stresses, as it simultaneously assesses all conditions and classes. We found that most co-accumulation modules contained peaks with high abundance in roots and low in leaves, or vice versa, again highlighting organs as primary drivers for metabolic diversity. One module (“cyan”) identified peaks specifically accumulated in Sym.Spore roots **(Supp. Fig. 19)**, a quarter of which were annotated as sphingolipids, again suggesting the importance of sphingolipids in AMS. Other modules contained peaks with more varied accumulation patterns. For example, the “turquoise” module identified peaks that were either specifically accumulated in hydroponics roots or excluded from them. The “gray60” module **(Supp. Fig. 19)** grouped peaks abundant in leaves but excluded from all roots except those experiencing AMS. These may represent foliar metabolites that undergo transport to the roots and play a role in symbiosis. A more detailed analysis of these peaks can reveal novel insights into the biochemistry of Brachypodium abiotic and biotic responses.

**Figure 5:**
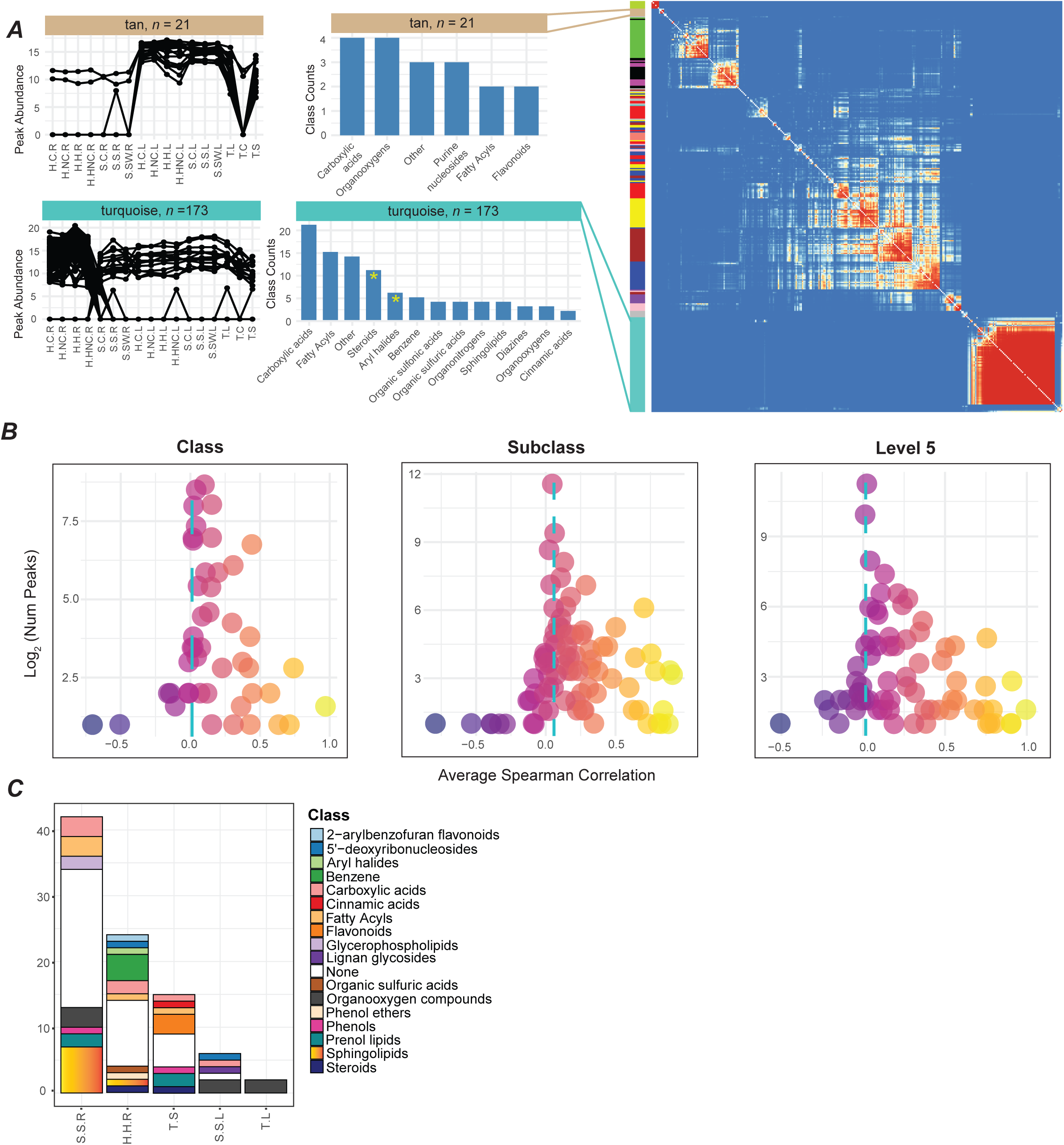
Characterizing Metabolite Co-abundance, Negative Mode. (A) WGCNA Topography Overlap Matrix, depicting correlations among peaks placed into significant modules. Normalized abundance patterns and CANOPUS Class distributions are plotted for selected modules (Class “None” not shown). An asterisk (*) denotes Classes that were significantly enriched in a module (Fisher’s exact test, FDR adjusted *p*-value < 0.05, count in module at least 5). Condition names in the abundance pattern plots (left) are abbreviated such that only the capital letters of the full names (seen in Figure 4) are shown. (B) Average pairwise Spearman correlation among peaks in the same CANOPUS Class, Subclass, or Level 5. The blue line shows average correlation among randomly selected peaks. (C) Counts of biomarkers found in each stress/tissue, colored by CANOPUS Class. Condition name abbreviations are as in (A).

A majority of WGCNA modules contained multiple Classes, and 7/18 modules were enriched in ≥1 Class (Fisher’s exact test, FDR adjusted p < 0.05). Some metabolite classes, such as Flavonoids, were enriched in multiple modules with differing abundance patterns (**Supp. Figs. 18, 19**). Cinnamic acids and flavonoids were usually significantly overrepresented in modules with higher accumulation in leaves than roots. Interestingly, cinnamic acids were perturbed substantially in leaves only under heat stress, while they were highly perturbed in roots under all conditions **(Supp. Fig. 16)**. Flavonoids, on the other hand, were significantly highly perturbed only in roots but not in leaves (**Supp. Figs. 16, 17**). These results point to differing regulation of individual metabolite classes in roots vs. leaves. Also, of the ten Classes enriched across all modules, in either positive or negative mode, seven were lipids, further highlighting their functional relevance.

To determine if “Class” is too broad a level for co-regulation, and if more evidence for co-regulation is found at the “Subclass” or “Level 5” level, the average pairwise Spearman correlation among peaks in the same Class, Subclass, or Level 5 category (**Fig. 5B** and **Supp. Fig. 17C**), was compared to the average correlation among randomly drawn peaks. At each hierarchy level, only a small number of classes had average correlation ≥0.5, and most classes had correlation close to random. Notably, at each level of the hierarchy, several classes were unusually large, with > 100 members, raising the possibility of low structural similarity within each class. Thus, we sought to determine whether class size and structural similarity within class contribute to average class correlation (see **Supplementary Methods** for calculation). Correlations between average class correlation and average class cosine score were consistently positive (**Supp. Figs. 20, 21**) suggesting greater structural similarity within a class translates to greater accumulation correlation. Correlations between class size and correlation/cosine score were negative, highlighting the importance of more specific class definitions. We note that overall, these metrics explained only a very low proportion of variance.

Taken together, these results indicate that while some classes (e.g. Flavonoids and their subclasses) may represent groups of co-regulated peaks, this is likely not the case for most classes at each level of the ontology. This may reflect the specificity of underlying metabolic and regulatory pathways, which may significantly increase concentrations of specific individual metabolites of a structural class. These results also suggest that utilizing the multi-label nature of the chemical ontology could be a better approach for finding peaks belonging to coordinated routes of metabolism rather than using single classes.

### Comparative analysis facilitates analysis of known metabolites and biomarker detection

Our dataset provides a unique opportunity to analyze the accumulation patterns of known metabolites, as well as find biomarkers, i.e. peaks that accumulate highly (not necessarily specifically; see **Supplementary Methods**) in one condition/organ. We selected salicylic acid (SA), abscisic acid (ABA), and naringenin for analysis as they were identified by GNPS with match scores ≥0.89 (ABA and naringenin were additionally correctly annotated by CANOPUS), and may be of relevance in the studied conditions. We further validated these identifications by uncovering their major fragments from the literature, and checking for matching fragments in our queries (**Supp. Table 3**). SA is known to accumulate in roots under AMS (Zhang et al., 2013) and, in some species, under heat (Hara et al., 2012). We found that SA accumulated (but not significantly increased) in AMS roots, and was mildly but significantly increased in Heat roots (t-test, p-value < 0.05) (**Supp. Fig. 22**). In contrast, ABA levels highly increased in AMS roots, and in Heat and HeatNoCopper leaves (t-test, p-values < 0.05). Finally, for naringenin, mean decreases were observed in roots for all conditions (significant decreases seen in Heat and AMS; t-test, p-values < 0.05) corroborating our observations of RDPI of the broader Flavonoid Class.

We also found that the numbers of biomarkers detected in each condition resembled the overall RDPI distribution --roots typically have more biomarkers than their foliar counterparts, and Spore roots and Heat roots have the highest numbers of biomarkers (**Fig. 5C** and **Supp. Fig. 17D; Supp. File 11**). We found 11-carboxyblumenol C glucoside to be a foliar biomarker for AMS, corroborating previously published data (Wang et al., 2018a) (**Supp. Fig. 23A**). We discovered other peaks that either share fragment peaks with the Blumenol C or exhibit fragment peaks of a Blumenol C core lacking a carboxyl group (**Supp. Fig. 23B, D**). We also detected a peak specific to Spore leaf and not present in AMS roots, which shares no fragment peaks with the Blumenol C and was classified by CANOPUS as a 5’-deoxyribonucleoside (**Supp. Fig. 23C**) --suggesting that AMS induces other foliar-specific routes of metabolism.

### Visualizing metabolite class importance using the BAR platform

In order to make the data described herein more easily accessible to the scientific community, these data were integrated into the Bio-Analytic Resource for Plant Biology (BAR) website as a novel electronic Fluorescent Pictograph (eFP) browser (available for testing at: https://bar.utoronto.ca/efp_brachypodium_metabolites/cgi-bin/efpWeb.cgi). CANOPUS Classes with at least five members in both positive *and* negative modes were included in this eFP browser. This eFP browser has two viewing options: with the Relative viewing option, the changes of metabolite Class levels across conditions can be readily observed (**Fig. 6**) as the average change in normalized peak area under a condition. With the Absolute viewing option, the average normalized peak areas are plotted per organ and condition. Besides showing how the Class changes in abundance across conditions, the Absolute view option also provides information about which ionization mode best illustrates changes experienced in that Class. Notably, for some Classes (e.g. Furanoid lignans, Purine nucleotides) we observe different changes in abundance across ionization modes. While this may be due to CANOPUS peak misannotations, especially for Positive mode, it may also reflect different subclasses being detected in different ionization modes. This finding has implications for targeted comparative metabolomics studies, as results obtained in one ionization mode may not necessarily hold in the other. By establishing our eFP browser, we seek to enable the community to draw further conclusions from our existing results, and facilitate the design of future comparative metabolomics and downstream validation studies.

**Figure 6:**
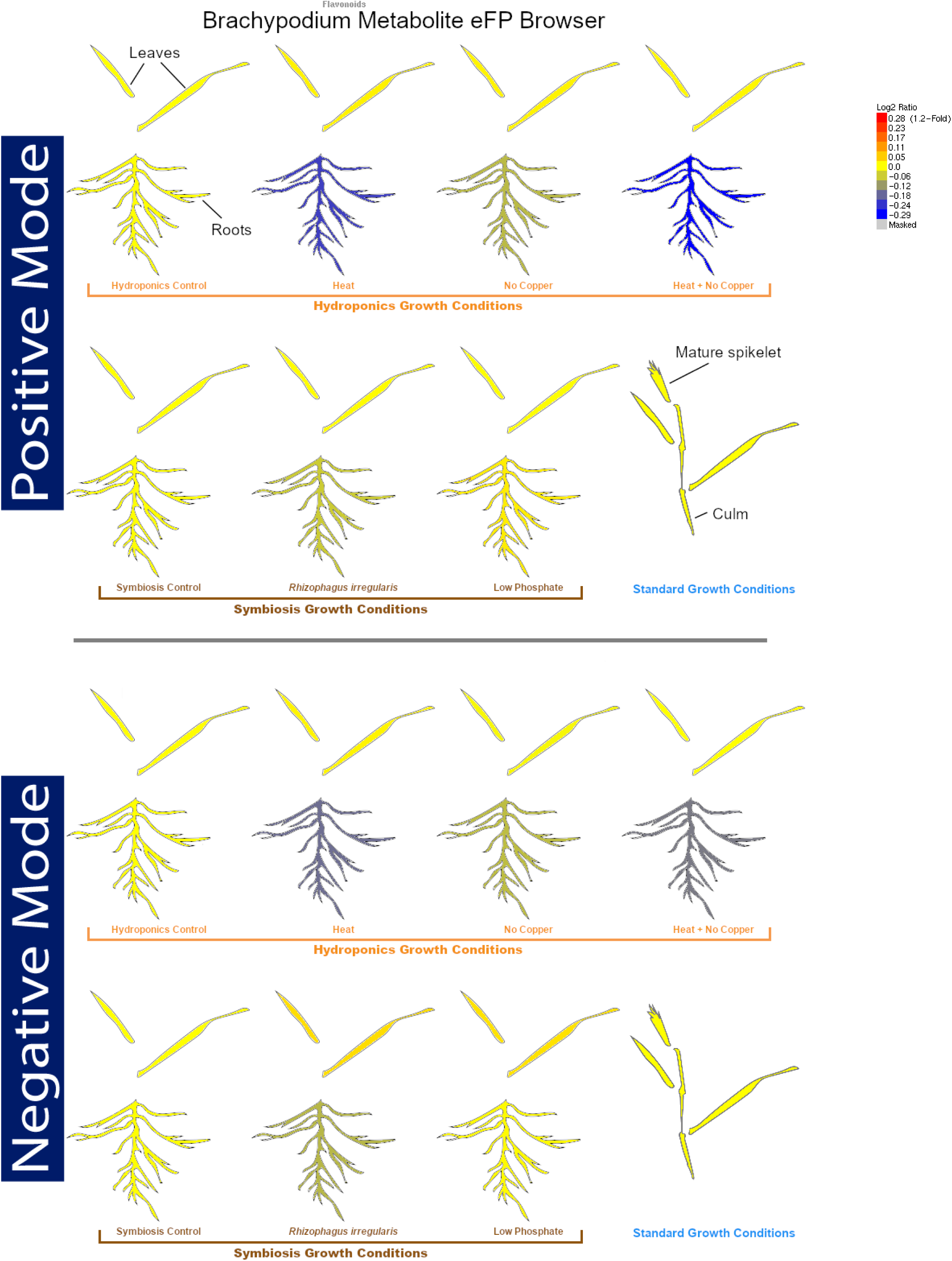
Visualizing Stress-induced Changes in Class Abundance. In Relative mode of the eFP browser (shown here for Flavonoids), the Log2 Fold Changes in average Class abundance are plotted between a condition and its control. The consistent decreases among stressed roots are again seen.

## Discussion

While recent improvements in LC-MS hardware have generated impressive advancements in metabolite detection, associating the thousands of metabolites detected in each species with biological processes remains an open challenge (Chaleckis et al., 2019). In this study, using three complementary approaches – information theory, ML-based analysis and co-accumulation clustering – of LC-MS data, we performed a more comprehensive analysis of metabolome perturbations of *B. distachyon* under different environmental conditions.

When applying information theoretic measures to the global metabolome, we found that roots are, on the whole, more stress-responsive than leaves, despite leaves having a more expansive and complex metabolome. The finding that leaves have consistently more peaks than roots may be due to biological or technical/processing reasons, as root harvesting required a washing/drying step to remove the attached soil particles, which may have also removed epidermal metabolites. While the increased number of peaks in foliar samples directly contributes to their increased diversity, the finding that leaf metabolomes are less perturbed under stress than roots is intriguing. Previous studies have also found roots to be more impacted than leaves under a variety of stresses, including heat (Giri et al., 2017), and salinity (López-Cristoffanini et al., 2021). Notably, drought stress --not included in our study --appears to be an exception in which leaves are more impacted than roots (Gargallo-Garriga et al., 2014), indicating that the greater metabolic plasticity of the roots is not universal. These results may again be due to technical considerations, as peaks with m/z > 800 were not detected, thereby excluding cuticular waxes, which are stress-responsive (Baker, 1974; Wang et al., 2018b; Kan et al., 2022). Additionally, highly polar and highly non-polar compounds were excluded from our data. Both roots and leaves contain such compounds, and therefore it is unclear how results would differ with these compounds included.

Our analyses revealed that the combined HeatNoCopper stress was less disruptive to the root metabolome than the Heat stress alone --suggesting that one week of Cu deficiency primed the roots for subsequent protection against heat stress response. Another interpretation is that critical heat response mechanisms were not activated in the roots after a week of Cu deficiency, which could therefore lead to decreased reproduction or long-term survival after heat. Since the recovery of these plants were not studied, it is not possible to ascertain which interpretation is correct. However, these results reveal an intriguing interplay between heat stress and Cu deficiency. In Arabidopsis, such an interplay is suggested through shared aspects of heat and Cu deficiency responses. For example, Cu deficiency triggers accumulation of ferric superoxide dismutase 1 to account for reduced activity of Cu/Zn superoxide dismutases (Abdel-Ghany and Pilon, 2008). This shift may help protect the roots against reactive oxygen species produced during later heat shock. Recent evidence has also suggested that SPL7, a master regulator of Cu deficiency (Yamasaki et al., 2009), may upregulate miR156 under Cu deficiency (Perea-García et al., 2021). In Arabidopsis, miR156 is induced after an initial heat stress event and provides heat shock memory, as plants lacking miR156 showed decreased growth and survival after subsequent heat events (Stief et al., 2014). As miR156 is also induced in wheat after heat stress (Xin et al., 2010), and as several miRNAs are known to have different induction patterns in different tissues (Sunkar et al., 2012), we hypothesize that miR156 upregulation under Cu deficiency helps prime Brachypodium roots for heat stress. Future molecular studies can help test these hypotheses.

In this study, we combined CANOPUS --a tool for structurally annotating peaks --with information theoretic and related measures to analyze more specific metabolome perturbations. Our study captured a multi-pronged, organ-differentiated metabolomic response of Brachypodium to environmental change comprising lipidomic perturbations and alterations of phenylpropanoid pathway products such as lignans, cinnamic acids, and flavonoids. Lipids, on the whole, are highly stress-responsive, with glycerolipids, GPs, sphingolipids and fatty acyls having high perturbations under several conditions. These perturbations may be a result of changes in membrane composition (known to occur under heat (Higashi and Saito, 2019) and low P stress (Nakamura, 2013)), and/or production of lipid signaling molecules, such as oxylipins (Ali and Baek, 2020) and sphingolipids (Berkey et al., 2012). Under AMS specifically, certain fatty acyls and GPs are known to be produced (Wewer et al., 2014; Bravo et al., 2017), and while this is indeed reflected in our data (**Supp. File 10**) we also found that other lipid Classes, such as Sphingolipids and Prenol lipids, were highly altered under AMS, suggesting that AMS has wide-reaching effects on the Brachypodium lipidome. We unexpectedly found that flavonoids are, on average, decreased in the roots in response to all conditions except low P – a finding corroborated by a focused assessment of naringenin. The training data for CANOPUS for flavonoids was also large (Dührkop et al., 2021) – given their high representation in public databases – thus, flavonoid class predictions are likely to be correct. This inference was also previously confirmed in sweet potato flavonoids and anthocyanins via comparison with MS/MS networking (Bennett et al., 2021). In general, flavonoids are known to be accumulated under several stresses (Ferdinando et al., 2012), yet the wholescale labelling of all flavonoids as antioxidants has been questioned (Agati et al., 2020). Several studies have additionally found disordered regulation of flavonoid biosynthesis, either at the level of individual flavonoids/flavonoid biosynthetic genes (Wu et al., 2020) or post-transcriptional regulation of flavonoid biosynthesis (Cui et al., 2019). These observations reveal a need for a deeper investigation of flavonoid roles and/or metabolic reprogramming under stress. We further found that WGCNA, a tool commonly used in RNA-seq studies, is effective at uncovering peaks with similar abundance patterns, which are potentially in the same routes of metabolism. Our study was also able to detect biomarkers, which can reveal novel insights into condition-specific activations of metabolic pathways.

In conclusion, we found that information theory metrics and chemical class predictions are effective tools to analyze comparative metabolomics data. Our results reveal a very dynamic plant metabolome influenced by multiple environmental and developmental factors. As more untargeted LC-MS/MS studies are performed, comparative analyses of these datasets may reveal common patterns and the core stress response across groups of plant species. The overall workflow described here can enable a more streamlined analyses of such untargeted datasets. Particularly, visualizing metabolomic data using the eFP browser may reveal hidden spatial differences in metabolome perturbations not easily discernible otherwise, and guide the design of targeted studies. For example, this visualization can be a useful tool to identify a better mode of ionization for molecules of interest as well as reveal metabolite classes to be assessed via targeted analyses. Our study shows that data-intensive analytical methods are useful for gleaning novel biological insights from untargeted metabolomics studies.

## Materials and Methods

### Plant Growth Conditions and Harvesting

*Brachypodium distachyon* Bd-21 seeds for plants used in the Symbiosis (Sym) and Tissue (Tis) experiments were sterilized in 10% (v/v) household bleach containing 0.005% (v/v) Tween-20 for seven minutes, thoroughly washed 5x in sterile water and germinated in petri dishes on moist Whatman filter paper in dark at 4 °C for seven days and three additional days at room temperature (RT). Germinated seedlings were incubated for additional 3-5 days under constant light while maintaining constant humidity. The germination protocol for plants used in the Hydro experiment was performed as outlined previously (Sheng et al., 2021). Additional details about plant growth conditions are described in **Supplementary Methods**. After the growth period, harvesting of all plant material took place between noon and 3pm to maintain circadian profiles of genes and metabolites. All samples were stored at −80 °C until further processing. We verified that the Cu-deficiency and AMS conditions worked as expected using RT-PCR of previously known condition-specific genes (Rahmati Ishka and Vatamaniuk, 2020) **(Supplementary Methods)**.

### Metabolite Extraction and Sample Preparation

All plant material was rough ground over liquid nitrogen using scissors to enable equal and homogenous separation for RNA and metabolite extraction. All samples were further subjected to bead homogenization using a mixer mill (Retsch, Haan, Germany) at 30 bpm with 1-minute intervals in 2 mL reaction tubes containing four 2.3 mm chrome steel beads. Ground samples were lyophilized overnight. Sample fresh weights (200 mg leaves, 550 mg roots, 150 mg spikelets and culms) were determined to ensure 50 mg of dry weight for all tissues. Samples were ground again in the bead homogenizer for 10 minutes, and centrifuged at 14000 g for 10 minutes in order to collect all powdered sample at the bottom of the tube. Metabolites were extracted using a mixture of acetonitrile, isopropanol, and water (ratio of 2:2:1) containing 0.1% (v/v) formic acid, and 30 uM of three internal standards (Telmisartan, Propyl-4-hydroxy-benzoate, and Kanamycin). After solvent addition, samples were vortexed several times over a period of 15 minutes to facilitate extraction. After centrifugation for 10 minutes at 16000 g to remove particulates, the samples were transferred into amber HPLC vials and stored at −80 °C until LC-MS analysis. Sample vials were shipped to the Joint Genome Institute on dry ice for LC-MS analysis, where LC-MS was performed using an Agilent 1290 Infinity LC system (Agilent, Santa Clara, CA) coupled to a Thermo QExactive HF orbitrap mass spectrometer (Thermo Scientific, San Jose, CA). Additional details are provided in the **Supplementary Methods**.

### Metabolomic data filtering, normalization, and imputation

All RAW files were converted to mzML format using ProteoWizard v 3.0.7230. TICs were made for all files of a given polarity using XCMS (Mahieu et al., 2016) (**Supp. Fig. 3**). All files of a given mode (positive or negative) were then imported into MS-DIAL v4.48 (Tsugawa et al., 2020) for peak deconvolution and alignment. Parameters files for positive and negative mode usage are supplied (**Supp. File 12**). The peak areas of the internal standards Telmisartan and Propyl-4-hydroxy-benzoate were manually checked to determine consistency across samples. For each polarity, MS-DIAL outputs a quantitative alignment file, displaying the peak areas of all metabolites in all samples, and a Mascot Generic Format (mgf) file of all fragmented metabolites. Detected metabolites were filtered, imputed, and normalized using a custom R script (developed in R v4.0.4) (R Core Team, 2020), available on GitHub (https://github.com/lizmahood/brachy_metabolomics) as described in **Supp. Fig. 4**. Metabolites eluting out at 90 seconds or earlier were removed as the Total Ion Current observed at the beginning of runs was high enough that accurate quantification of metabolite values could not be assured (**Supp. Fig. 3**). Imputation was performed with the R package impute and VSN was performed with the R package vsn (Huber et al., 2002). Normalization was performed using NOREVA (Li et al., 2016a), followed by identification of differentially accumulated metabolites, both of which are described in more detail in **Supplementary Methods**.

### Peak annotation with CANOPUS

The mgf format MS-DIAL output files were filtered to remove adducts and peaks detected in Blank samples using an in-house python script (https://github.com/lizmahood/brachy_metabolomics). The CANOPUS module (Dührkop et al., 2021), included in the SIRIUS4 v4.9.8 software suite (Dührkop et al., 2019) was used to annotate singly charged peaks with their probable structural classes, as defined in the multilabel ChemOnt ontology (Djoumbou Feunang et al., 2016). The Zodiac module (Ludwig et al., 2020) was additionally used to improve each peak’s predicted molecular formula (which CANOPUS uses for annotation). For each compound, CANOPUS predicts the “Parent Class” – the class of the largest substructure in the molecule – and outputs the probability that the predicted Parent Class is correct, based upon its training data. Other predictions are made at different hierarchies of the ontology (Superclass, Subclass, etc). Any annotation with prediction probability < 0.5 was not considered in downstream analyses. Additionally, if a classification was discarded for not meeting this probability threshold, each subsequent prediction (at more specific hierarchies) was removed as well, regardless of their prediction probabilities.

### Peak Identification with GNPS and MSDIAL

The “All Public MS/MS” msp files provided by MSDIAL (http://prime.psc.riken.jp/compms/msdial/main.html#MSP) were used for identification. To remove false positive identifications, we imposed a threshold of >0.8 for both the Dot Product and Reverse Dot Product scores between the query and database match. Feature based molecular networking through GNPS (Nothias et al., 2020) workflow v28.2 was additionally used for peak identification. Spectral database libraries included those publicly available in GNPS, as well as the NIST 17 library, which was kindly provided by JGI. All parameters for molecular networking were kept at default values excepting: Precursor ion mass tolerance - 0.01 Da, Library search min matched peaks - 3, Top results to report per query - 20, Score threshold - 0.4, Maximum analog search mass difference - 200. We again imposed a threshold of >0.8 for the match score between the query and database match, and only considered the top 1 match per query.

To compare annotations between peak identifications and CANOPUS, InChIs of identified compounds were converted to InChI-Keys through the chembl_ikey python module, and structural classifications were obtained with ClassyFire Batch (https://cfb.fiehnlab.ucdavis.edu/).

### MS/MS molecular networking

MS-FINDER v3.44 (Tsugawa et al., 2016) was used to perform molecular networking using the filtered mgf files, with the following parameters: Mass tolerance 0.01, Relative abundance cutoff 5%, MS/MS similarity cutoff 70%, RT tolerance 100. The Superclass of each peak, as well as the conditions each peak was identified as Differentially Abundant in, were added to the node file. The edge file and this augmented node file were imported into Cytoscape v.3.8.0 (Su et al., 2014) for figure generation using the Prefuse Force Directed Layout.

### Estimating information theoretic measures

The following information theoretic metrics were calculated for our dataset as described previously (Li et al., 2020): Hj (the Shannon entropy/Metabolomic profile diversity), Si (Metabolomic specificity), and δj (Metabolome specialization index). The Relative Distance Plasticity Index (RDPI), as calculated for all peaks in each stress condition, was also determined as described previously (Valladares et al., 2006). The RDPI calculation was applied to the entire metabolome, and then applied separately to each compound class (for compound classes with at least five peaks classified into them). The RDPI formula was amended in order to determine if a class is up- or down-accumulating under a stress. Briefly, for each condition-control pair of samples, a distribution of the abundance changes of all peaks was made, and the mean change in peak abundance was calculated per class. Let *d*_*ij*→*i′j*′_ represent the peak area changes to all peaks *i* common to a condition-control pair (*j*→*j′*). The mean value of the peak area change for each compound class was computed as ∑(*dij* → d*i*′*j*′) / *n*, where *n* is the number of peaks per class. For each condition, these per-class mean values were compared to the overall distribution of *d*_*ij*→*i′j′*_ across all metabolite peaks to determine the percentile of the per-class value with respect to the peak area changes of all compared metabolites. For the purposes of plotting in Fig. 4, the classes with percentiles >70 (large average increase in abundance) or <30 (large average decrease in abundance) and at least five members were identified, and up to five classes with the highest/lowest percentiles were plotted.

### Weighted Correlation Network Analysis Construction and Module Analysis

Using the Weighted Correlation Network Analysis (WGCNA) R package (Langfelder and Horvath, 2008), an unsigned adjacency network was made from the normalized area of all fragmented peaks. The soft powers were 129 and 131 in positive mode and negative mode, as these were the lowest values achieving a R^2^ of at least 0.8. Hierarchical clustering via the hclust function was performed using method = “average”. The minimum module size was 10. All peaks that failed to be assigned to a module were discarded, and the remaining peaks were re-clustered into a dendrogram, and visualized alongside their Topography Overlap Matrix. The CANOPUS class of all peaks in each significant module was determined. Each class (except “None”) was analyzed for enrichment in a particular module if there were at least 5 members in the module. Enrichment was calculated using a Fisher’s exact test with all fragmented peaks as the background population.

### Visualizing CANOPUS Class Abundance on the BAR Platform

Briefly, an input image was generated representing the experiments described in this paper. The eFP Browser code (Winter et al., 2007) was then modified in several ways to be able to display CANOPUS data. First, the color scheme was modified from the default yellow-red color scheme of the original eFP Browser, to make a visual distinction between the metabolite data being displayed in the modified version and transcript data displayed in the original browser. Second, because CANOPUS data have a lower dispersion, we introduced a possibility of setting a minimum value for the color scale other than zero. Last, CANOPUS classes with at least five members in both Positive *and* Negative ionization modes were included in this eFP browser, and were databased in such a way that the data from the two modes could be retrieved separately. CANOPUS data may be freely explored at https://bar.utoronto.ca/efp_brachypodium_metabolites/cgi-bin/efpWeb.cgi.

## Data availability

The LC-MS/MS data is deposited on the GNPS website with the accession ID MSV000089340. All code developed for analyses is available on the GitHub repository (https://github.com/lizmahood/brachy_metabolomics) also forked on the moghelab GitHub page.

## Author contributions

Conceived the study: GDM, MJH, OKV, NJP

Planned experiments: Authors that conceived the study plus EHM, AAB, LHK, AB, MRI, YJ, VL

Performed experiments: EHM, AAB, KK, LHK, BPB, KBL, AB, MRI, YJ, VL

Wrote the manuscript: EHM, NJP, GDM

Reviewed the manuscript: All authors

## Acknowledgements

GDM and EHM would like to thank the Cornell BioHPC Center for assisting with computing infrastructure and the Cornell and Boyce Thompson Institute Greenhouse staff for growth chamber maintenance. GDM would like to thank Dr. Trent Northen for initial discussions during project development. EHM would like to thank Drs. Kai Dührkop and Sebastian Böcker for explanation of CANOPUS outputs and Dr. Dapeng Li for clarification of the RDPI formula.

## Funding sources

This research was funded by Cornell Startup Funds and US DoE-Joint Genome Institute grant #504788 to GDM, USDA-NIFA grant #2021-67034-35227 to EHM, Deutsche Forschungsgesellschaft award #411255989 to LHK, US DOE BER grant #DE– SC0012460 to MJH, USDA-NIFA grants #2018-67013-27418 and #2021-67013-33798 to OKV, and NSERC and the Genome Canada/Ontario Genomics OGI-128 to NJP. The work 10.46936/10.25585/60001229 conducted by the U.S. Department of Energy Joint Genome Institute (https://ror.org/04xm1d337), a DOE Office of Science User Facility, is supported by the Office of Science of the U.S. Department of Energy operated under Contract No. DE-AC02-05CH11231.

## Conflicts of Interest

No conflicts of interest exist.

## Notes

### Competing Interest Statement

The authors have declared no competing interest.

