## Supplemental Figures for "Characterization and visualization of global metabolomic responses of *Brachypodium distachyon* to environmental changes"

*Supplemental Figure 1*

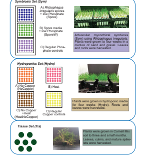


***Supplementary Figure 1: Experimental design and set-up of the three sub-experiments studied.*** The growth stages and tissues harvested in each set are shown in the figure. Names in parentheses refer to condition names used in the text. Although each condition is shown as a separate block for illustrative purposes, for Sym, the position of the plants in each set was randomized in the growth chamber, and for Hydro, the position of hydroponics containers was randomized. In the Symbiosis Set (referenced as Sym in the text), plants were grown in one of three conditions. Plants in the "Spore" set received 1mL of *Rhizophaguz irregulariz* spores and low phosphate (20 uM) fertilization, plants in the "SporeW" set received 1mL of fungal growth media lacking spores and low phosphate fertilization, and plants in the "Control" set received regular phosphate (200 uM) fertilization. In the Hydroponics set (Hydro in the text), plants in the "NoCopper" set received one week of media lacking Copper after three weeks of media with standard Copper amounts. Plants in the "Heat" set received one day of 40 C after four weeks of growth. Plants in the "HeatNoCopper" set received both stresses, while plants in the "Control" set received neither.

***Supplementary Figure 2***

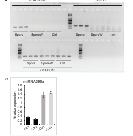


***Supplementary Figure 2: sq-PCR and RT-qPCR for Validation of Cu and AMS Treatments.*** (A) Roots from three biological replicates (each a pool of 20 plants) per condition (Spore = inoculation with *Rhizophagus irregularis* (*Ri*), SporeW = inoculation with fungal media lacking spores) were assessed for expression of symbiosis marker genes. *Bd*UBC18 was used as a control. The number of cycles were optimized to ensure the total cDNA concentration remained in the exponential phase. (B) Roots from two biological replicates (each a pool of 12 plants) from Hydro.Control and NoCopper roots were used to quantify the expression of miR398a, a Copper deficiency marker, using RT-qPCR. *Bd*Actin was used as the reference gene for data normalization. Error bars depict the Standerd Error of measurements across three technical replicates.

*Supplementary Figure 3*

*A*

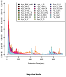


*B*

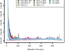


***Supplementary Figure 3: Total Ion Chromatograms for all samples.*** (A) Positive mode, and (B) Negative mode. "BLK" = Blank, "CP" = NoCopper, "CTRL" = Control, "H" = Heat, "HCP" = HeatNoCopper, "S" = Spore, "SW" = SporeW, "Spklt" = Spikelet, "L" = Leaf, "R" = Root. Due to the large spike in intensity, peaks with Retention Time < 90 seconds were removed.

***Supplementary Figure 4***

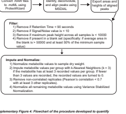


***Supplementary Figure 4: Flowchart of the procedure developed to quantify metabolites.*** For more details, see Methods and Supplemental Methods.

***Supplementary Figure 5***

A

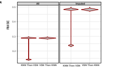


B

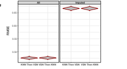


***Supplementary Figure 5: Root Mean Squared Error (RMSE) between true values and imputed values under two imputation-normalization orders.*** (A) positive and (B) negative mode. "All" quantifies the RMSE between all true normalized metabolite peak areas and all normalized peak areas. "Imputed" quantifies the error between *only* the imputed values. The smaller error values occurring in positive mode in "KNN Then VSN" is attributed to the VSN formula, and did not influence our decision to impute before normalizing.

*Supplementary Figure 6*

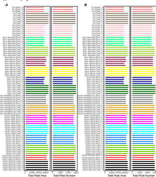


***Supplementary Figure 6: Total peak area and peak numbers per sample.*** Total peak counts were determined after peak filtration and imputation for (A) positive and (B) negative mode. Total peak areas were determined after peak filtration, imputation, and normalization to the dry sample weight and using VSN. Samples that were deemed outliers based on correlation or PCAs were removed from downstream analyses and are not shown (three samples were removed from Tis.Spike, one from Sym.Ctrl.Root, and two from Hydro.NoCop.Root).

*Supplemetary Figure 7*

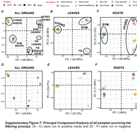


***Supplementary Figure 7: Principal Component Analysis of all samples surviving the filtering process.*** (A - C) were run in positive mode and (D - F) were run in negative mode. (A) and (D) depict all organs, (B) and (E) include only leaf samples, and (C) and (F) show only root samples. The color in each PCA corresponds to the condition, and the shape to the organ. The ellipses and annotations addded to the positive mode PCAs are for aiding interpretation only and have no further meaning.

***Supplementary Figure 8***

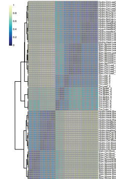

***Supplementary Figure 8: Replicate Dendograms.*** All non-oulier positive samples were clustered using a distance matrix of 1 - pairwise Pearson's correlation. Samples were clustered based on filtered and normalized metabolite values.

**Supplementary Figure 9**

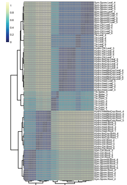


***Supplementary Figure 9: Replicate Dendrograms.*** All non-outlier negative samples were clustered using a distance matrix of 1 - pairwise Pearson's correlation. Samples were clustered based on filtered and normalized metabolite values.

***Supplementary Figure 10***

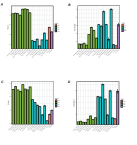

***Supplementary Figure 10: Diversity and Specialization per condition.*** Bars represent the mean of each metric per condition, and error bars show the standard deviation across all replicates remaining after outlier removal. (A) and (B) show the Diversity and Specialization in positive mode, while (C) and (D) show them in negative mode.

**Supplementary Figure 11**

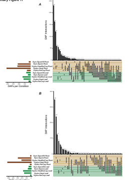

***Supplementary Figure 11: Full UpSet plots showing intersections of Differentially Accumulated Peaks (DAPs) in all sets of conditions.*** DAPs include both up- and down- accumulated peaks. (A) includes peaks detected in positive mode and (B) includes negative mode. The vertical bar charts (showing DAP Intersections) show the numbers of DAPs found in each intersection of conditions/organs, as outlined by the combination of dots underneath the bar.

***Supplementary Figure 12***

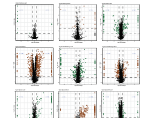

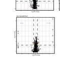

***Supplementary Figure 12: Volcano Plots of Differentially Accumulated Peaks in Negative Mode.*** q-values are the Benjamani-Hochberg corrected p-values resulting from an unpaired t-test. The cutoffs for significant fold changes were determined as outlined in Methods. Large, colored dots represent peaks with statistically significant abundance differences (DAPs). Dots with a log2 fold change of 2.5 or -2.5 were not present in the control or stress, respectively.

**Supplementary Figure 14**

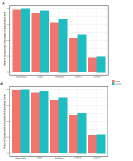


***Supplementary Figure 14: Ratio of Compounds Annotated at Different Levels of the ChemOnt Ontology*** (A) Positive mode and (B) Negative mode. Nonfiltered bars represent all compound classifications. Filtered bars represent only those classifications with posterior probabilities > 0.5.

**Supplementary Figure 15**

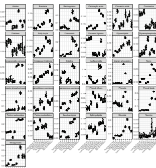


***Supplementary Figure 15: RDPI per CANOPUS Class, Positive Mode.*** Each dot represents the RDPI between one replicate in a particular stress and its control. "None" represents metabolites that received no CANOPUS Class-level annotation.

***Supplementary Figure 16***

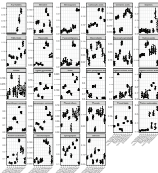


***Supplementary Figure 16: RDPI per CANOPUS Class, Negative Mode.*** Each dot represents the RDPI between one replicate in a particular stress and its control. "None" represents metabolites that received no CANOPUS Class-level annotation.

### Supplemental Figure 17

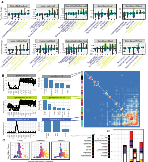

**_Supplemental Figure 17: Charting Stress-Induced Shifts of Molecular Classes._** (A) Abundance changes of peaks in response to stress. Each stress depicts Classes that were the most accumulated (Class name in yellow) or diminished (Class name in purple), on average. For a Class to be plotted, its average value must be greater than the 70th percentile or lower than the 30th percentile of all stress-induced peak area changes. Individual metabolites are plotted as circles, outliers are shown as +. (B) WGCNA Topography Overlap Matrix, depicting correlations among peaks placed into significant modules. Normalized abundance patterns and CANOPUS Class distributions are plotted for selected modules (Class "None" not shown). An asterisk (*) denotes Classes that were significantly enriched in a module (Fisher's exact test, FDR adjusted p-value < 0.05, count in module at least 5). Condition names in the abundance pattern plots (left) are abbreviated such that only the capital letters of the full names (seen in A) are shown. (C) Average pairwise Spearman correlation among peaks in the same CANOPUS Class, Subclass, or Level 5. The blue line shows average correlation among randomly selected peaks. (D) Counts of biomarkers found in each stress/tissue, colored by CANOPUS Class. Condition name abbreviations are as in (B).

### Supplementary Figure 18

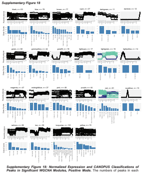

***Supplementary Figure 18: Normalized Expression and CANOPUS Classifications of Peaks in Significant WGCNA Modules, Positive Mode.*** The numbers of peaks in each module is shown. Peaks receiving no CANOPUS classifications (classification of "None") not shown. Classes with fewer than 3 peaks in a module were deemed "Other". An asterisk (*) denotes Classes that were significantly enriched in a module (Fisher's exact test, FDR adjusted *p*-value < 0.05, count in module at least 5). Condition names in the expression plots are as in Figure 4, section B. Peak abundances in the red and lightgreen modules are colored so as to differentiate distinct expression patterns.

*Supplementary Figure 19*

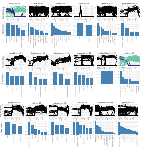


***Supplementary Figure 19: Normalized Expression and CANOPUS Classifications of Peaks in Significant WGCNA Modules, Negative Mode.*** The numbers of peaks in each module is shown. Peaks receiving no CANOPUS classifications (classification of "None") not shown. Classes with fewer than 3 peaks in a module were deemed "Other". An asterisk (*) denotes Classes that were significantly enriched in a module (Fisher's exact test, FDR adjusted *p*-value < 0.05, count in module at least 5). Condition names in the expression plots are as in Figure 4, section B. Peak abundances in the black and pink modules are colored so as to differentiate distinct abundance patterns.

**_Supplementary Figure 20_**

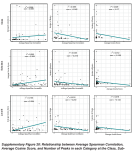

***Supplementary Figure 20: Relationship between Average Spearman Correlation, Average Cosine Score, and Number of Peaks in each Category at the Class, Subclass, and Level 5 levels.*** All plots are for peaks in Positive mode. The "corr" on each plot is the Pearson correlation between the variable pair.

**Supplementary Figure 21**

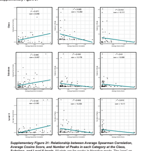


***Supplementary Figure 21: Relationship between Average Spearman Correlation, Average Cosine Score, and Number of Peaks in each Category at the Class, Subclass, and Level 5 levels***. All plots are for peaks in Negative mode. The "corr" on each plot is the Pearson correlation between the variable pair.

***Supplementary Figure 22***

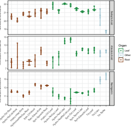


***Supplementary Figure 22: Normalized Abundance Values for Salicylic Acid, Abscisic Acid, and Naringenin Across All Organs and Conditions.*** Metabolites were identified through a database search (using GNPS, see Methods).

*Supplemental Figure 23*

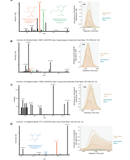


***Supplemental Figure 23: Substructures of Foliar Biomarkers of Symbiosis.*** (A) 11-carboxyblumenol C 9-O-glucoside was identified to Level 2. Characteristic "Blumenol" fragment peaks are shown in green and orange. (B) Unknown compound with no accumulation in Sym Spore roots yet containing two characteristic "Blumenol" fragment peaks. (C) Unknown compound lacking any "Blumenol" fragment peaks. (D) Unknown compound with one "Blumenol" fragment peak and another fragment not observed in (A) yet likely representing a Blumenol substructure. Asterisks signify foliar symbiosis biomarkers, and each compound's ID corresponds with its Alignment ID in Supplementary File 3.
