## Supplemental Methods for "Characterization and visualization of global metabolomic responses of *Brachypodium distachyon* to environmental changes"

### Supplementary Methods

#### *Plant Growth Conditions and Harvesting*

For the symbiosis experiment, plants were grown for four weeks in a sterilized soil mixture of 2:2:1 play sand: coarse black sand: gravel as previously described (Floss et al., 2017). These plants were potted in 20.5 cm cones containing a large marble at the bottom to facilitate water movement and inhibit the release of soil when watering. Four plants were planted per cone, and cone placement within the growth chamber was randomized across conditions. Plants were inoculated with surface sterilized *Rhizophagus irregularis* spores by adding 1 mL of water containing 1000 spores/mL. Spores were placed onto a sand layer 7 cm below the soil surface. A control group (SporeW) received 1 mL of the media used to wash fungal spores to control for effects of fungal exudates. Plants were watered daily using a pump mist bottle and were fertilized weekly using a variant of Hoagland's solution (Javot et al., 2011). All components have ¼ strength except for phosphate (Potassium Phosphate monobasic), which was kept at 20 uM final concentration for the Spore and SporeW groups. For the phosphate control plants, 200 uM phosphate was used. Symbiosis and tissue plants were grown in a chamber with a 16/8 hours light/dark rhythm, ~155 µmol photons $m^{-2}$ $s^{-1}$ light intensity, 50% relative humidity, and 25 °C. Tissue plants were grown in Cornell Mix (Boodley and Sheldrake, 1972) for 3 months (to ensure development of spikelets) with five plants per pot. These plants were fertilized once weekly with 200 uM phosphate ¼ Hoagland's solution.

Hydroponics plants were grown in plastic containers containing 4.5 L of hydroponics media equipped with a floating Styrofoam bed holding the plants in small holes. 30 plants were grown in each container. Plants in the Control and Heat groups received hydroponics media throughout the whole experiment as described previously (Sheng et al., 2021) containing 250 nM Cu, whereas plants in the NoCopper and HeatNoCopper groups received hydroponics media containing no Cu for the final seven days of their growth. Hydroponics plants were grown in their own growth chamber, with the following growth conditions: 150 umol photons $m^{-2}$ $s^{-1}$, 50% relative humidity, with a 16/8 hours light/dark rhythm, and 24 °C day/18 °C night. Plants in the Control and NoCopper groups were grown for 28 days, and plants in the Heat and Copper-Heat groups were grown for an additional, final day at 40 °C.

Harvesting of all plant material took place between noon and 3pm to maintain circadian profiles of genes and metabolites. All tissues were placed on dry ice and flash frozen in liquid nitrogen after harvesting. For Symbiosis plants, five replicates were included for each group (Control, Spore, and SporeW), with five cones (20 plants) harvested per replicate, and roots and leaves harvested from each cone. For Tissue plants, six replicates were harvested, with five plants per replicate, and leaves, culms and spikelets harvested from each plant. For the Hydroponics plants, five replicates were harvested per group (Control, Heat, NoCopper, and HeatNoCopper), with 12 plants per replicate, and leaves and roots harvested per plant. To minimize batch effects, each replicate contained plants collected across two containers. All samples were stored at -80 °C until further processing.

***RNA extraction and validation of positive control genes***

RT-qPCR analysis was performed to ensure the efficacy of the Copper deficiency treatment, while semi quantitative RT-PCR was done to detect mycorrhizal growth in Spore roots. RNA was initially extracted as follows. Using approximately 75mg of root tissue, RNA was pre-extracted using Trizol® (Invitrogen, Life Technologies) according to the manufacturer's instructions, but with two additional phase separation steps involving the addition of 200 and 100 µL chloroform, respectively, to the aqueous phase and subsequent centrifugation. The Monarch® Total RNA Miniprep Kit (NEB, Ipswich, MA) protocol was followed for extraction of total RNA from the final aqueous layer and the RNA was eluted in 50 uL nuclease-free water. For cDNA synthesis, 500 ng RNA was reverse transcribed using the Protoscript II Reverse Transcriptase (New England Biolabs, Ipswich, MA [NEB]), with Oligo-dT17 primer (**Supp. Table 3**), according to manufacturer recommendations at 45 ºC for one hour. After heat inactivation for 20 minutes at 80 ºC, the reaction mix was diluted with four volumes of nuclease-free water and stored at -20 ºC until further use.

For semi-quantitative PCR the Q5 Hot Start High Fidelity DNA Polymerase (NEB) with dNTPs (200 uM each) and primers (0.5 µM each, **Supp. Table 3**) were used. To ensure sampling occurred before the amount of PCR product reached saturation, aliquots were taken after 25, 30, and 35 cycles, with 30 cycles ultimately used as the products remained in the exponential phase (**Supp. Fig. 2**). The levels of the UBC18 transcript were used for normalizing between different samples. Products were analyzed on a 2% (w/v) agarose gel stained with 0.01% (v/v) EtBr. Three biological replicates (each a pool of 20 plants) were used from the Spore, SporeW and Sym.Control treatments. For Copper deficiency, RT-qPCR was performed with the Copper deficiency marker miR398 as described previously (Rahmati Ishka and Vatamaniuk, 2020), using the primers listed in **Supp. Table 3**. Two biological replicates (each a pool of twelve plants) were used from the Copper deficiency and Hydro.Control treatments. RT-qPCR analysis was conducted using iQ SYBRGreen Supermix (Bio-Rad) according to manufacturer's instructions in the CFX96 real-time PCR system (Bio-Rad).

***Ultra-High Performance Liquid Chromatography (UHPLC) – Tandem mass-spectrometry (MS/MS) Run Conditions***

In preparation for LC-MS analysis, 1.5 uL of 100 ug/mL internal standard (2-Amino-3-bromo-5-methylbenzoic acid, Sigma) in 100% methanol was added to 150 uL of sample extract in 2:2:1 isopropanol:acetonitrile:water with 0.1% formic acid, followed by a brief vortex. Liquid chromatography mass spectrometry (LC-MS) analysis was performed using an Agilent 1290 Infinity LC system (Agilent, Santa Clara, CA) coupled to a Thermo

QExactive HF orbitrap mass spectrometer (Thermo Scientific, San Jose, CA). For each sample, 2 uL were injected into a reverse phase C18 column (Agilent ZORBAX Eclipse Plus C18, Rapid Resolution HD, 2.1 x 50 mm, 1.8 µm) held at 60 ºC at a flow rate of 0.4 mL/min. The column was equilibrated with 100% buffer A (100% water w/ 10 mM ammonium formate, adjusted to pH=3.2 using formic acid) for 1 minute, followed by a linear gradient to 100% buffer B (95:5 acetonitrile:water w/ 10 mM ammonium formate) over 11 minutes, isocratic elution for 1.5 minutes, then returning to 100% buffer A over 1 minute then 1.5 minutes isocratic elution. Full MS spectra were collected from *m/z* 100-800 at 60,000 resolution in both positive and negative ion mode, in centroid format, with MS/MS fragmentation data acquired using stepped then averaged collision energies of 10-20-40 eV, or 20-50-60 eV, for the top 10 MS1 ions per scan at 17,500 resolution. The upper m/z limit was set at a relatively low value in order to maximize quantification accuracy. Mass spectrometer source settings included a sheath gas flow rate of 55 (au), auxiliary gas flow of 20 (au), sweep gas flow of 2 (au), spray voltage of 3 kV and capillary temperature of 400 ºC. Sample injection order was randomized, with an injection blank of 100% methanol run between each sample, as well as a quality control (QC) mix run every 20 samples consisting of a mixture of 3 sample extracts.

***Normalization methods comparison with NOREVA***

The normalization scheme used was chosen after comparing different normalization methods on our data using the web application NOREVA (Li et al., 2016), two of which were based on Internal Standards and three were "data-driven". The normalization schemes chosen were: VSN, Cross-contribution Compensating Multiple standard Normalization (CCMN) (Redestig et al., 2009), Normalization through Optimal selection of Multiple Internal Standards (NOMIS) (Sysi-Aho et al., 2007), and Probabilistic Quotient Normalization (PQN) with Power Scaling. NOMIS and CCMN normalize using two of our internal standards (Telmisartan and Propyl-4-hydroxy benzoate), while the other methods normalize across all metabolites. The data loaded into NOREVA was pre-filtered, imputed, and removed of outlier samples prior to upload, following the pipeline in **Supp. Fig. 4** (concluding just before the VSN step). Additionally, all metabolite values of 0 were changed to 0.01 before upload, to preempt NOREVA's mandatory imputation step. No data removal or transformations or were performed by NOREVA prior to normalization.

***Determination of imputation and normalization order***

To determine if imputation followed by normalization, or vice versa, introduced more error into the dataset, we compared imputed values made by both orders to actual metabolite values following the protocol of (Shah et al., 2017). Briefly, the percentage of zeros across the entire dataset was quantified, then all metabolites with a 0 in any sample were removed. "Missing" values were introduced into this "complete" dataset by randomly turning values to 0, such that the percentage of "missing" values was equal to the original percentage of zeros in the entire dataset. Next, imputation followed by normalization, and normalization followed by imputation were performed, and the resulting metabolite values were compared to the true values. Error was quantified by Root Mean Squared Error.

#### ***Metabolite differential accumulation analysis***

As VSN is non-linear, applying it to a dataset will drastically change the fold changes for some metabolites, e.g. metabolites with non-normalized fold changes well above 2 may have their fold changes decrease substantially after normalization, to well below 2. Due to this, when determining differentially abundant metabolites, we considered their fold changes before and after normalization. Additionally, a metabolite had to have a Wilcoxon Rank-Sum p-value < 0.1 and a Benjamini-Hochberg adjusted p-value < 0.05 (resulting from an unpaired t-test) to be considered differentially abundant in a particular condition.

Metabolites with a non-normalized fold change of ≥2 were considered as potentially differentially abundant. To determine fold change cutoffs for normalized differential abundance, we used a stepwise approach to account for the non-linear nature of VSN normalization. First, the distribution of normalized fold changes of all metabolites failing to meet the non-normalized fold change cutoffs was determined. The 99$^{th}$ and 1$^{st}$ percentiles of this distribution were used as the cutoffs for normalized fold change (specifically: 0.2662 and -0.2727 in positive mode, and 0.2919 and -0.2681 in negative mode). The extreme percentiles were chosen so as to minimize the number of metabolites considered with a non-normalized fold change ≤2. To be considered differentially abundant, metabolites must have met both normalized and non-normalized fold change cutoffs, and have both fold changes be in the same direction (e.g. a metabolite with a significant increase in a condition before normalization must retain the increase after normalization). Thus, our definition of differentially abundant metabolites is very stringent.

#### ***Pairwise Cosine Score Calculation***

Before Cosine score calculation, peaks were centroided such that all peaks with mass differences < 0.01 Da were represented by the most abundant peak only. The pairwise cosine score between two peaks was calculated as a Neutral Dot Product (NDP) as follows (Li et al., 2020):

$$\frac{\left(\sum_i^{S1\&S2} W_{S1,i} W_{S2,i}\right)^2}{\sum_i W_{S1,i}^2 \, W_{S2,i}^2}$$

where S1 and S2 denote, respectively, spectrum 1 and spectrum 2, and $W_{S1,i}$ and $W_{S2,i}$ indicate peak intensity-based weights. The numerator of the equation considers the common peaks differing by less than 0.01 Da among the two spectra (peak intersections), whereas the denominator considers all peaks in each spectrum (peak union). Weights were calculated as:

$$W = [\text{Peak Intensity}]^{m}\,[\text{Mass}]^{n}$$

where $m = 0.5$ and $n = 2$ as used previously (Horai et al., 2010; Li et al., 2020).

***Biomarker Identification***

Steps for identifying biomarkers for a condition are as follows: (i) for each peak, find the median peak area per condition, (ii) for each peak, find $x = i / j$, where $i$ is the maximum median peak area, and $j$ is the second highest median peak area, (iii) find the fold-change threshold, the 95$^{th}$ percentile of all $x$. These values were 1.235 and 1.416 in positive mode and negative mode, respectively. (iv) Peaks with $x$ greater than the fold change threshold, and with low abundance (normalized abundance < 14) in all other conditions, were considered putative biomarkers. All such peaks were manually checked for high-quality peak shapes.

**Boodley JW, Sheldrake R** (1972) Cornell Peat-Lite Mixes For Commercial Plant Growing. Information Bulletin 43, New York State College of Agriculture and Life Sciences, Cornell University, Ithaca, NY 8

**Floss DS, Gomez SK, Park H-J, MacLean AM, Müller LM, Bhattarai KK, Lévesque-Tremblay V, Maldonado-Mendoza IE, Harrison MJ** (2017) A Transcriptional Program for Arbuscule Degeneration during AM Symbiosis Is Regulated by MYB1. Current Biology **27**: 1206–1212

**Horai H, Arita M, Kanaya S, Nihei Y, Ikeda T, Suwa K, Ojima Y, Tanaka K, Tanaka S, Aoshima K, et al** (2010) MassBank: a public repository for sharing mass spectral data for life sciences. Journal of Mass Spectrometry **45**: 703–714

**Javot H, Penmetsa RV, Breuillin F, Bhattarai KK, Noar RD, Gomez SK, Zhang Q, Cook DR, Harrison MJ** (2011) Medicago truncatula mtpt4 mutants reveal a role for nitrogen in the regulation of arbuscule degeneration in arbuscular mycorrhizal symbiosis. The Plant Journal **68**: 954–965

**Li B, Tang J, Yang Q, Cui X, Li S, Chen S, Cao Q, Xue W, Chen N, Zhu F** (2016) Performance Evaluation and Online Realization of Data-driven Normalization Methods Used in LC/MS based Untargeted Metabolomics Analysis. Sci Rep **6**: 38881

**Li D, Halitschke R, Baldwin IT, Gaquerel E** (2020) Information theory tests critical predictions of plant defense theory for specialized metabolism. Science Advances **6**: eaaz0381

**Rahmati Ishka M, Vatamaniuk OK** (2020) Copper deficiency alters shoot architecture and reduces fertility of both gynoecium and androecium in Arabidopsis thaliana. Plant Direct **4**: e00288

**Redestig H, Fukushima A, Stenlund H, Moritz T, Arita M, Saito K, Kusano M** (2009) Compensation for Systematic Cross-Contribution Improves Normalization of Mass Spectrometry Based Metabolomics Data. Anal Chem **81**: 7974–7980

**Shah JS, Rai SN, DeFilippis AP, Hill BG, Bhatnagar A, Brock GN** (2017) Distribution based nearest neighbor imputation for truncated high dimensional data with applications to pre-clinical and clinical metabolomics studies. BMC Bioinformatics **18**: 114

**Sheng H, Jiang Y, Rahmati M, Chia J-C, Dokuchayeva T, Kavulych Y, Zavodna T-O, Mendoza PN, Huang R, Smieshka LM, et al** (2021) YSL3-mediated copper distribution is required for fertility, seed size and protein accumulation in Brachypodium. Plant Physiology **186**: 655–676

**Sysi-Aho M, Katajamaa M, Yetukuri L, Orešič M** (2007) Normalization method for metabolomics data using optimal selection of multiple internal standards. BMC Bioinformatics **8**: 93
